# Horn trophy seized in illegal wildlife trade assigned to swamp buffalo using a combined morphometric and DNA based approach in wildlife forensics in India

**DOI:** 10.1101/2021.09.27.461958

**Authors:** Vipin, Vinita Sharma, Chandra Prakash Sharma, Surendra Prakash Goyal, Sandeep Kumar Gupta

## Abstract

The illegal wildlife trade has threatened the existence of many extant wild animal species throughout the world. While dealing with the illegal wildlife trade of horns, we face problems of not having a proper protocol and lack of reference database to assign the species for proper implementation of wildlife laws. In one such condition, a horn trophy suspected to be of a wild buffalo was seized by authorities and sent to us for species identification. We used a combined approach of morphological and DNA analysis to ascertain the seized horn’s species. The two measurements, circumference at the base (CAB) and length on the front curve (LOFC) were measured for the seized and other horns of different bovid species, showing morphological resemblance with the seized horn. The 3-D scatter plot, generated by the values of CAB, LOFC and CAB/ LOFC, differentiated the different bovid species into distinct clusters and placed the seized horn in the proximity of domestic buffaloes. The Bayesian evolutionary analysis of the partial D-loop gene (521bp) placed the seized horn in a clade with swamp buffaloes. Since swamp buffaloes are domestic buffaloes, both these approaches concluded the same results. Hence, the current protocol developed may also be used to differentiate among wild buffalo, domestic buffalo, Cattle, Wild yak, Gaur and Takin using a combined approach of morphometric and DNA-based analysis, which may be used to deal with illegal wildlife trade of different bovid species at the world level.

## Introduction

Illegal wildlife trade is a major threat to many extant wild animal species of the world. To combat the illicit wildlife trade, the species-level identification of seized wildlife parts or their products is required for the legal prosecution of accused/s in the court of law (1). The bovids formed the next most abundant group of mammals after carnivores in the illegal trade market (2). Almost all wild bovid species are in trade worldwide (3-5) serve as big and small game animals for their magnificent horn trophies, head mounts, skin/fur, bones, meat and other body parts. Horns, which is the characteristics feature of species of family Bovidae, makes these species as important big game throughout the world from time immemorial and also serve as a key ingredient of traditional medicines worldwide from time immemorial till date (2, 6, 7). Hunting trophies are an important part of growing global commerce and almost all of the species of hoofed mammals are prized as hunting trophies (3-5).

In many instances, the lack of proper species identification protocols hinders the implementation of wildlife protection laws at local and international levels. In one such condition, a horn suspected to be of wild buffalo was seized by the law enforcement agencies in India and was sent to Wildlife Institute of India, Dehradun, to identify the species of its origin. As no defined protocol is available to identify the species of a horn, we used a combined approach of morphometric and DNA-based techniques for its species identification. This kind of combined approach has been proved to be a very promising tool while identifying the species of fake tiger claws in the illegal wildlife trade (8). The use of a combined approach always has the edge over using a single one as one may come to rescue when the other has failed or both may complement each other. As mentioned above, the horns are the characteristics feature of species belonging to the family Bovidae; 19 species of wild bovids are enlisted in either schedule of Wildlife (Protection) Act, 1972 of India. These species’ intact horns are chiefly confiscated in a huge amount at various checkpoints by the enforcement agencies. Apart from wild, the comparison of the seized horn with domestic bovid species also makes it essential here for leaving no doubt about excluding any horn type of bovid family. The lack of a comparative morphological and genetic reference database for a wide range of bovid horns makes it challenging to identify the species of a horn or its parts and products.

In the present study, we generated and collected the morphometric data of the horns of wild and domestic bovid species. The partial D-loop genetic data is available for almost all domestic buffalos of India, South-East Asia, the Mediterranean and China in the published literature (9). Therefore, we generated the partial D-loop gene data for the seized horn in the present study.

Hence, we tried to prove how a combined approach of morphometric and DNA based approach can be utilized to identify the species of a horn trophy seized in the illegal wildlife trade. The researchers may use the protocol developed here to identify all three species of buffalos (wild, river and swamp) along with some other wild bivide species in the world.

## Material and methods

### Case history and sample collection

We used a horn trophy with metal coverings on its tip and at the base, suspected to be of a wild buffalo. It was seized by the authorities in Delhi, India, in April 2014 and sent to the Wildlife Institute of India, Dehradun, for species identification (Fig. 1).

**Figure 1.**
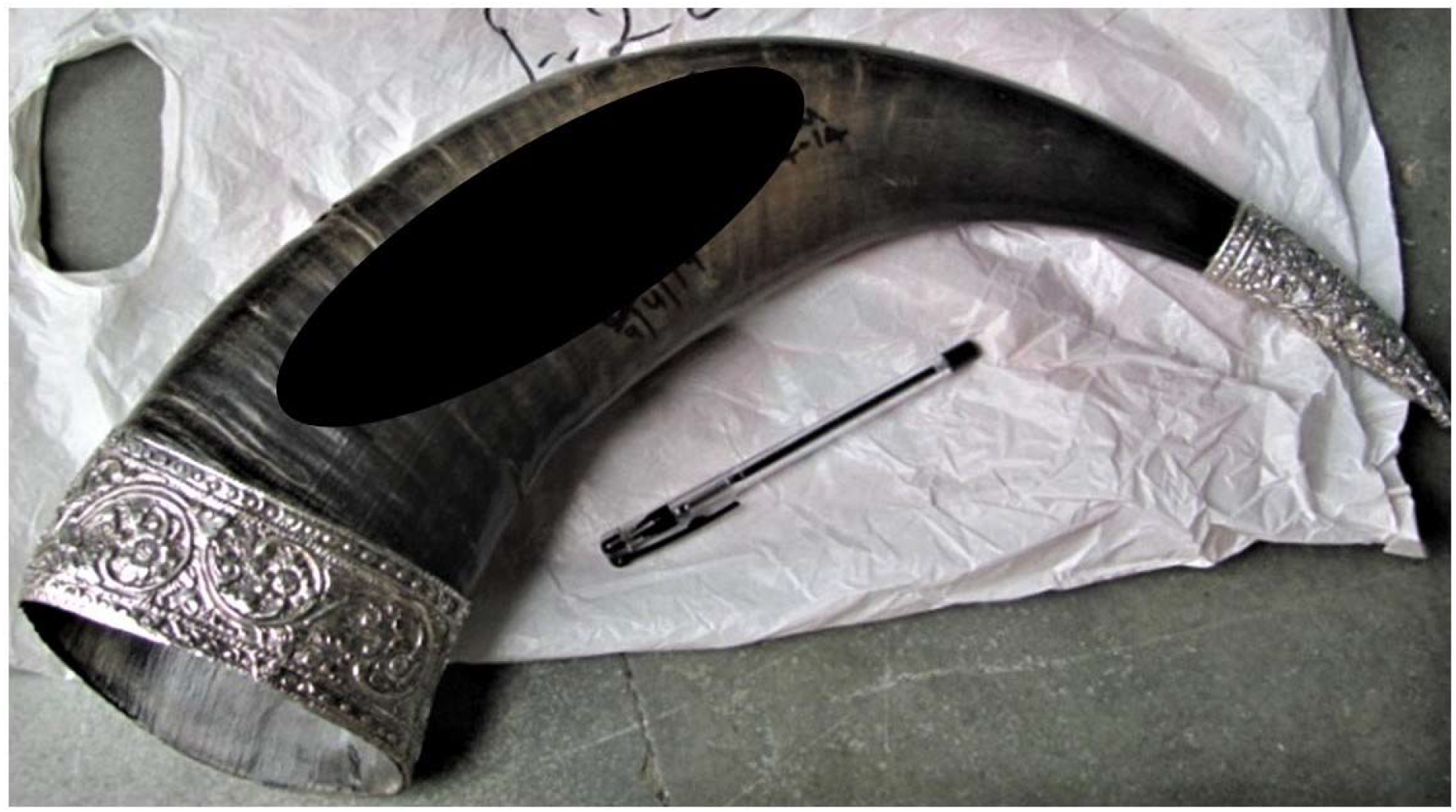
Seized horn trophy.

### Morphometric analysis

A burn test was performed to check the genuineness of the biological originality of the confiscated samples for the presence of keratin as per Vipin et al. (8). The available morphological and morphometric characters were noted and compared with the horn characteristics of Wild buffalo and other species which were showing resemblance to the seized horn (Wild water buffalo, *Bubalus arnee* Kerr, 1792; Wild yak, *Bos mutus* Przewalski, 1883; Gaur, *Bos gaurus* Smith, 1827; domestic Asian water buffalo, *Bubalus bubalis* Lydekker, 1913 (river + swamp type). The length on the front curve (LOFC) and circumference at the base (CAB) of the seized horn were measured and for other bovid species mentioned above, were taken from the published literature (10-14) as well as self-collected (Sharma et al., *unpublished*) and are given in Table 1 (Note-only the minimum and maximum values are shown here, the complete dataset is available on request). A 3-D scatter plot was generated using LOFC and CAB in SPSS statistics software version 19 (15).

**Table 1.** Details of horn measurements for the comparison of morphometric analysis.

| Species | N |  | LOFC (in cm) | CAB (in cm) | LOFC/CAB |
| --- | --- | --- | --- | --- | --- |
| Wild Water Buffalo | 37 | Minimum | 126.37 | 30.00 | 0.22 |
|  |  | Maximum | 196.53 | 58.42 | 0.43 |
| Wild Yak | 24 | Minimum | 76.20 | 34.29 | 0.41 |
|  |  | Maximum | 97.16 | 43.18 | 0.52 |
| Gaur | 43 | Minimum | 53.00 | 40.00 | 0.53 |
|  |  | Maximum | 88.27 | 54.61 | 0.85 |
| Gayal | 5 | Minimum | 20.00 | 24.42 | 1.02 |
|  |  | Maximum | 23.96 | 28.39 | 1.42 |
| River Buffalo | 16 | Minimum | 41.00 | 15.37 | 0.25 |
|  |  | Maximum | 80.36 | 32.00 | 0.78 |
| Swamp Buffalo | 25 | Minimum | 31.56 | 19.93 | 0.46 |
|  |  | Maximum | 63.09 | 30.90 | 0.80 |
| Seized Horn | 1 | Value | 61.00 | 36.00 | 0.59 |

### DNA analysis

#### Genomic DNA extraction

First of all, the small flakes were scrapped from the base of the horn’s inner surface with the help of a sterile scalpel and discarded since the outer layer might have contaminants and interfere with PCR amplification. Then the flakes were scrapped a second time and collected in a sterile Petri dish. 500 mg of these scrapped flakes of the horn were used for the extraction of the genomic DNA using a modified protocol developed by Rohland and Hofreiter (16), which uses silica binding in the presence of a high concentration of guanidium thiocyanate. The protocol was modified by incubating the sample in extraction solution at 56□ in a water bath with agitation for 36 hours. The DNA elution was done using 100 µl Elution buffer from DNeasy blood and tissue kit (Qiagen, Hilden, Germany). All steps of DNA extraction were carried out in a sterile and dedicated laboratory to avoid any contamination. One negative control was also included to cross-check any contamination due to reagents.

#### Species identification and phylogenetic analysis

D-loop region of the mitochondrial genome was amplified using a primer pair (F: 5’TAGTGCTAATACCAACGGCC3’; R: 5’AGGCATTTTCAGTGCCTTGC 3’) described by Sutarno (17). The conditions for polymerase chain reaction and DNA sequencing were followed as described by Kathiravan *et al* (18) and Vipin *et al* (19), respectively. The sequence generated from D-loop gene was used in a BLAST search for the identification of species. Published sequences for D-loop gene (9) of domestic (river and swamp type) water buffalos from Bhadwari (AF475159), Toda (AF475264), South Kanara (JQ394841), Murrah (AF475204), Marathwada (JQ394848), Chilika (JQ394858), Surti (AF475249), Pandharpuri (AF475234), Jaffarabadi (AF475174), Nagpuri (AF475222, AF475219), Mehsana (AF475189), Lower Assamese (JQ394874, JQ394876), Upper Assamese (JQ394852, JQ394851, JQ394854, JQ394853), Manipur (JQ394867, JQ394866, JQ394869), South-East Asia (AF197219), Mediterranean (AF197199) and Chinese (DQ364160, DQ364161) were included in the analysis to examine the phylogeny of the aligned sequences for the D-loop gene. The sequences of cattle (AB085926; 20), takin (FJ006534; Wu et al; Unpublished) and yalk (AY374125; Zhao et al; Unpublished) were also included in the study.

The Bayesian evolutionary analysis was done using BEAST2 v2.6.2 (21). The HKY substitution model was selected with Gamma Category Count 4. The empirical frequencies were selected for getting a good fit to the data. The strict molecular clock model was selected for the data. The Yule Model of speciation was selected for the tree prior as it has been considered more appropriate when using different species sequences. The priors for rest parameters were set as default. The MCMC chain length was set to 10 million for this data set. The trace log and tree log frequencies were set to 1000 and the screen log frequencies were set to 10000. Assessment of adequate sampling, other of BEAST2 v2.6.2 output parameters and run-length were done with TRACER (22). The tree file generated by BEAST2 v2.6.2 having 10000 trees was visualized on the DensiTree2 (23). The DNA sequence identity matrix was generated through Multiple Sequence Alignment using MUSCLE (24).

## Results

Upon preliminary investigation and the burn test performed, it was observed that the seized horn was made up of an outer keratinous sheath and is a bovid horn. The 3-D scatter plot separated each bovid species, used here, into a distinct cluster (Fig. 2). The seized horn was found in the proximity of the domestic buffalo group (river and swamp type) (Fig. 2). However, the species of the seized horn was still not clear from the scatter plot. Therefore, the seized horn was further subjected to the phylogenetic analysis to know the species of its origin. The comparison of the outer shape and measurements of the seized horn with other species completely ruled out its Wild water buffalo origin as was thought earlier by the authorities. It also becomes clear from the above analysis that these two measurements i.e., CAB and LOFC have the power to differentiate the horns of bovid species used in this study.

**Figure 2.**
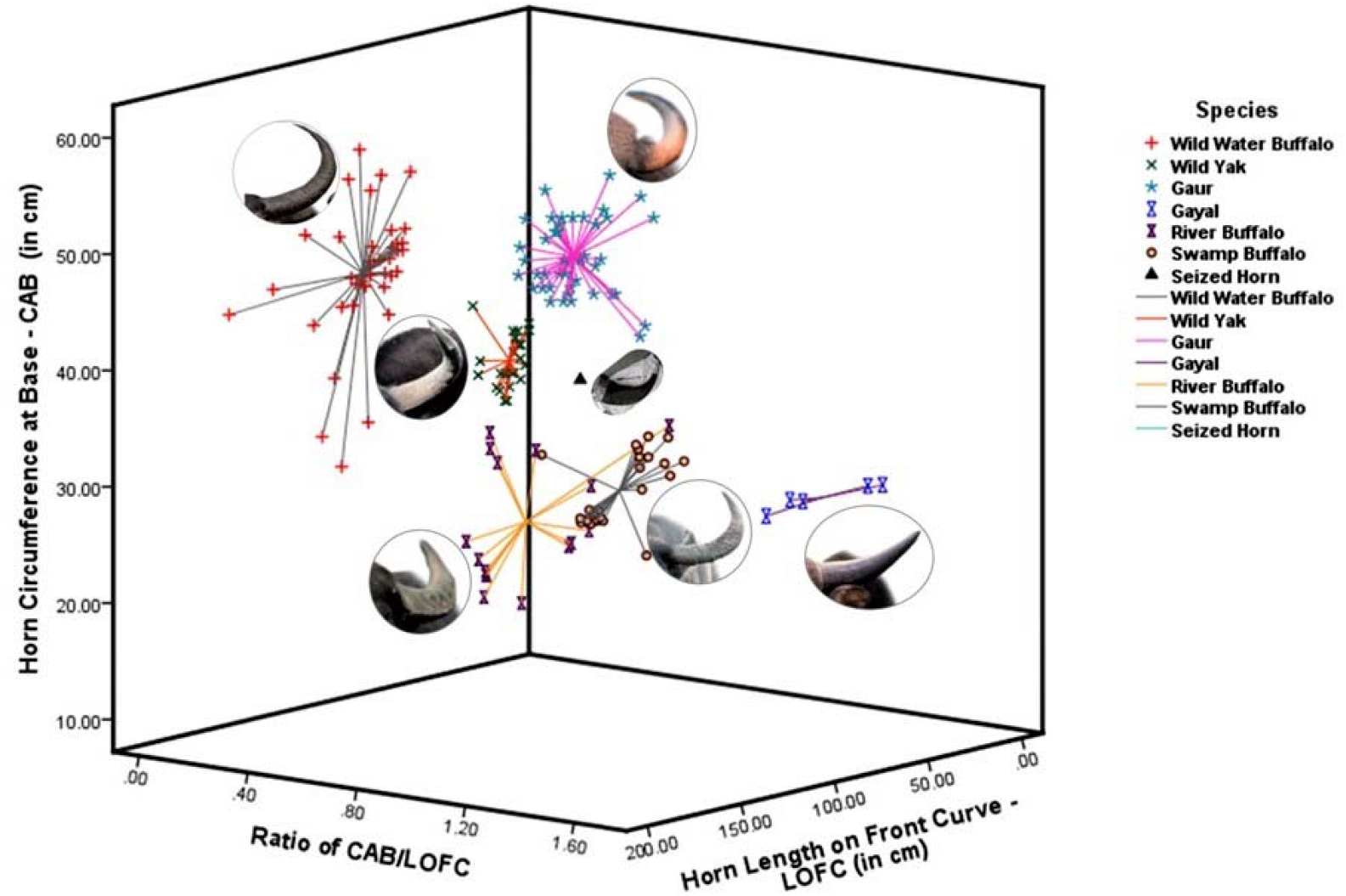
3D Scatter Plot drawn between the horn measurements.

Good quality of DNA was extracted from the horn trophy and successfully amplified by PCR using primers for the D-loop gene. This is for the first time that the protocol originally developed for the extraction of ancient DNA from bones and teeth by Rohland and Hofreiter (16) was modified and used for the extraction of DNA from a horn. The generated DNA sequence was used for NCBI (GenBank) BLAST, which indicated that the sequence belonged to *Bubalus bubalis*. The DNA sequence of the seized horn trophy was aligned with other 28 bovid species for the D-loop gene (9) and trimmed to make consensus sequences of 521 bases. The maximum clade credibility tree constructed using the D-loop gene sequences using BEAST2 v2.6.2 (21) separated the river and swamp buffalo populations into distinct clades. It showed that the seized horn trophy was placed in the clade constructed mostly of the Chinese and Southeast Asian swamp buffaloes (Fig. 3). The seized horn showed 99.42% - 99.62% D-loop DNA sequence similarity with Chinese and Southeast Asian Swamp buffaloes; however, with Manipuri Swamp buffalos, it was 98.43 – 99.41% (Table 2). Thus, the above results indicated that the seized horn was belonging to the Swamp buffaloes.

**Figure 3.**
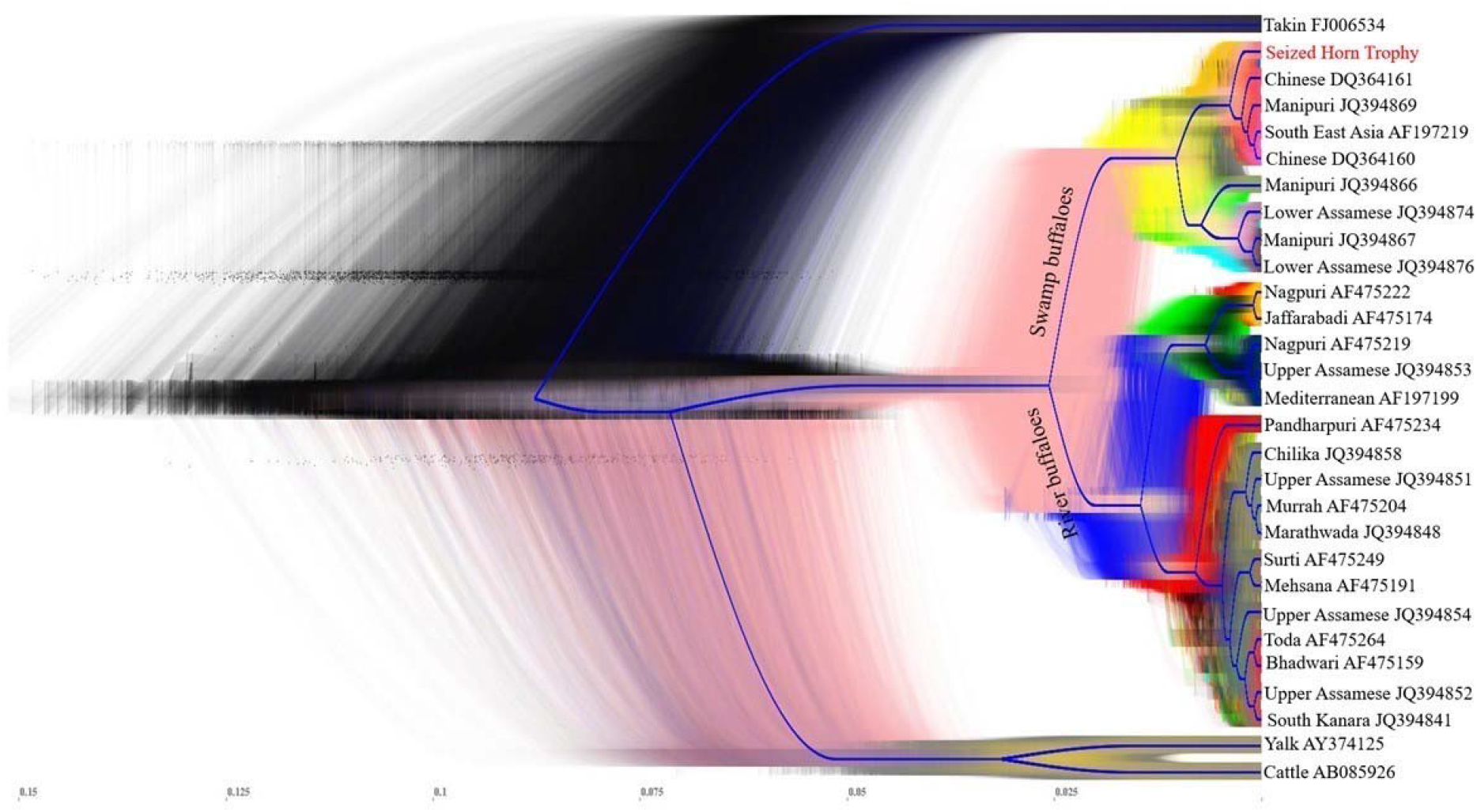
All phylogenetic trees with consensus tree (blue colour) showing seized horn trophy position with swamp buffalos.

**Table 2.** The D-loop sequence similarity percentage of seized horn trophy with other buffalo breeds and species.

| S. No. | Buffalo breed/ species | Sequence similarity with<br>seized horn trophy in<br>percent |
| --- | --- | --- |
| 1 | Seized Horn Trophy | 100.00 |
| 2 | Chinese DQ364160 | 99.62 |
| 3 | Southeast Asia AF197219 | 99.62 |
| 4 | Chinese DQ364161 | 99.42 |
| 5 | Manipuri JQ394869 | 99.41 |
| 6 | Manipuri JQ394866 | 98.63 |
| 7 | Manipuri JQ394867 | 98.43 |
| 8 | Lower Assamese JQ394874 | 98.43 |
| 9 | Lower Assamese JQ394876 | 98.43 |
| 10 | Upper Assamese JQ394853 | 96.27 |
| 11 | Pandharpuri AF475234 | 96.16 |
| 12 | Nagpuri AF475219 | 96.16 |
| 13 | Mediterranean AF197199 | 96.15 |
| 14 | South Kanara JQ394841 | 95.88 |
| 15 | Upper Assamese JQ394854 | 95.87 |
| 16 | Upper Assamese JQ394852 | 95.87 |
| 17 | Surti AF475249 | 95.78 |
| 18 | Bhadwari AF475159 | 95.78 |
| 19 | Toda AF475264 | 95.78 |
| 20 | Marathwada JQ394848 | 95.69 |
| 21 | Upper Assamese JQ394851 | 95.68 |
| 22 | Mehsana AF475191 | 95.59 |
| 23 | Murrah AF475204 | 95.59 |
| 24 | Jaffarabadi AF475174 | 95.59 |
| 25 | Nagpuri AF475222 | 95.59 |
| 26 | Chilika JQ394858 | 95.48 |
| 27 | Yalk AY374125 | 89.67 |
| 28 | Cattle AB085926 | 88.93 |
| 29 | Takin FJ006534 | 87.40 |

## Discussion

Hunting for the horn trophy has threatened the wild populations of buffalos and has also been responsible for the elimination of other longhorn animals from the gene pool (25). In past both the intense and selective harvesting of many animal species have induced evolutionary changes in behavior and phenotype (26-28). The selective hunting, preferably for bighorn size, had even more severe repercussions when it leads to reduced horn length with irreversible genetic values (28). Because of certain reasons the morphological appearance of swamp buffalo horn is more close to the wild type (http://factsanddetails.com/asian/cat62/sub408/entry-2830.html), easy access of their horns in comparison to wild types, most probably as a result of increased surveillance in protected areas by the authorities, the chances are more of swamp buffalo horn being traded easily, knowingly or unknowingly, in the name of wild buffalo horns to fetch more price in illegal wildlife trade. Though we used D-loop gene fragment to assign species of seized horn trophy, we could not compare it with the wild buffalo gene sequence because of the only one D-loop sequence available on NCBI (AF016397) with 14.8% comparable bases to our used sequences in this study and having only four nucleotide difference with swamp type buffalo. Though our results showed more than 99% D-loop DNA sequence similarity of the seized horn with Chinese and Southeast Asian Swamp buffaloes, the geographic population assignment is not easy unless the data of the complete gene is compared. Moreover, unlike wild animal populations, region-specific signatures are not expected in domestic animal populations since they are being traded from one place to another. Hence, taking here of geographic population assignment, in this case, seems irrelevant.

Morphologically, we analyzed the outer shape and morphometric measurements of bovid species and tried to identify the species. We played with the many horn measurements but the CAB and LOFC showed the robust characteristics to differentiate between different bovid species horns. Moving ahead, we tried the same measurements on nineteen wild bovid species of India and all these species were differentiated into distinct clusters in a 3-D scatter plot (Sharma et al.; Unpublished). So, for the first time in the world, these measurements have shown the ability to differentiated between wild as well as domestic bovid species and can be applied to other species also. The present combined methodology is complementing and validating each other as Wild water buffalo species was ruled out here by morphological analysis and common species (swamp buffalo, river buffalo and yalk) were differentiated by both techniques. The use of a combined technique is also recommended when either of the technique may not work because of certain reasons.

The current study also provides a unique protocol to differentiate between Wild and domestic buffalo, Cattle, Wild yak, Gaur and Takin using a combined approach of morphometric and DNA analysis. Which may be used to deal with illegal wildlife trade of different bovid species at world level.

## Acknowledgments

This study was funded by a grant-in-aid of the WII from the Ministry of Environment, Forest and Climate Change, Government of India.We gratefully acknowledge the support of the Director and Dean, WII, for this study.

## Declarations

### Conflicts of interest

There is no conflicts of interests.

### Availability of data and material

Not applicable.

### Code availability

Not applicable

### Ethics approval

Not applicable.

## Notes

### Competing Interest Statement

The authors have declared no competing interest.

## References

1. PAW (Partnership for Action against Wildlife Crime) (2005): Wildlife crime: a guide to the use of forensic and specialist techniques in the investigation of wildlife crime. Department for Environment, Food and Rural Affairs, Bristol.

2. Whiting MJ, Williams VL and Hibbitts TJ (2011) Animals traded for traditional medicine at the Faraday market in South Africa: species diversity and conservation implications. Journal of Zoology. 284: 84–96.

3. CITES (2003) CITES Identification Guide - Hunting Trophies. www.ucipfg.com/Repositorio/ELAP/cites_s/recursos/Trophy_Guide_HR.pdf.

4. Mallon D (2013) Trophy hunting of CITES – listed species in Central asia. Published by CITES Secretariat, Geneva, Switzerland. https://cites.unia.es/cites/file.php/1/files/cb-trophy-hunting.pdf.

5. IFAW (2016) Killing for Trophies: An analysis of global trophy hunting trade. http://www.ifaw.org/sites/default/files/IFAW_TrophyHuntingReport_UK_v2.pdf

6. Coghlan ML, Haile J, Houston J, Murray DC, White NE, Moolhuijzen P, Bellgard MI and Bunce M (2012) Deep Sequencing of Plant and Animal DNA Contained within Traditional Chinese Medicines Reveals Legality Issues and Health Safety Concerns. PLoS Genet 8(4): e1002657. doi:10.1371/journal.pgen.1002657

7. Chen J, Jiang Z, Li C, Ping X, Cui S, Tang S, Chu H and Liu B (2015) Identification of ungulates used in a traditional Chinese medicine with DNA barcoding technology. Ecology and Evolution. 5(9):1818–1825.

8. Vipin, Vinita Sharma, Chandra Prakash Sharma, Ved P. Kumar and Surendra Prakash Goyal (2016) Pioneer identification of fake tiger claws using morphometric and DNA based analysis in wildlife forensics in India. Forensic Science International. 266, 226–233. doi:10.1016/j.forsciint.2016.05.024.

9. Mishra BP, PK Dubey, B Prakash, P Kathiravan, S Goyal, D K Sadana, G.C. Das RN, Goswami Bhasin, BK Joshi and R S Kataria (2015) Genetic analysis of river, swamp and hybrid buffaloes of north-east India throw new light on phylogeography of water buffalo (Bubalus bubalis). Anim. Breed. Genet. 132: 454–466 doi:10.1111/jbg.12141

10. Ward R (1903) Records of Big Game. 4th Edition. Rowland Ward Limited, London.

11. Rasali DP, Joshi HD, Patel RK and Harding AH (1998b) Phenotypic clusters and karyotypes of indigenous buffaloes in the Western Hills of Nepal. Lumle Agricultural Research Station, Pokhara, Nepal. Technical Paper No. 98/2: 24 pp

12. Berthouly C, Rognon X, Van T Nhu, Berthouly A, Hoang H Thanh, Bed’Hom B, Laloe D, Chi C Vu, Verrier E and Maillard JC (2010) Genetic and morphometric characterization of a local Vietnamese Swamp Buffalo population. Journal of Anim. Breeding and Genetics.127. 74–84.

13. Kalita, R., Dandapatli, A., Choudhury, K.B.D., Das, G.C. and R.N. Goswami (2010): Conformation Traits of Swamp Buffalo of Assam at Different Age Groups. Indian J. Anim. Res., 44 (4) : 300–302.

14. Gubbawar S G, Shelke RR, Chavan SD and Pohare SR (2012) Phenotypic characteristics of gaolao strain of Nagpuri buffalo breed. The Asian Journal of Animal Sciences. Volume 7 Issue 1. 6–14.

15. IBM Corp, IBM SPSS Statistics for Windows, Version 19.0, IBM Corp., Armonk, NY, 2010.

16. Rohland Nadin and Hofreiter Michael (2007) Ancient DNA extraction from bones and teeth. Nature Protocols. Vol.2 No.7, 1756–1762

17. Sutarno Lymbery AJ (1997) New RFLPs in the mitochondrial genome of cattle. Anim. Genet., 28, 240–241

18. Kathiravan P, Kataria RS, Mishra BP, Dubey PK, Sadana DK, Joshi BK (2011) Population structure and phylogeography of Toda buffaloes throw light on possible origin of aboriginal Toda tribe of South India. J. Anim. Breed. Genet., 128, 295–304.

19. Vipin, Sharma Vinita and Gupta S K (2017) Molecular identification of victim species and its sex from the ash: a case of burning alive leopard (Panthera pardus). International Journal of Legal Medicine. DOI 10.1007/s00414-017-1619-1. Published online: 06 June, 2017

20. Fujise H, Murakami M, Devkota B, Dhakal IP, Takeda K, Hanada H, Fujitani H, Sasaki M and Kobayashi K (2003) Breeding distribution and maternal genetic lineages in Lulu, a dwarf cattle population in Nepal. Journal Anim. Sci. J. 74, 1-5 (2003)

21. Bouckaert R, Vaughan T G, Barido-Sottani J, Duchêne S, Fourment M, Gavryushkina A, Joseph Heled, Graham Jones, Denise Kühnert, Nicola De Maio, Michael Matschiner, Fábio K. Mendes Nicola F. Müller, Huw A. Ogilvie, Louis du Plessis, Alex Popinga, Andrew Rambaut, David Rasmussen, Igor Siveroni, Marc A. Suchard, Chieh-Hsi Wu, Dong Xie, Chi Zhang, Tanja Stadler, Alexei J. Drummond (2019) BEAST 2.5: An advanced software platform for Bayesian evolutionary analysis. PLoS computational biology. 2019; 15(4), e1006650.

22. Rambaut A, Drummond AJ, Xie D, Baele G and Suchard MA (2018) Posterior summarisation in Bayesian phylogenetics using Tracer 1.7. Systematic Biology. syy032. doi:10.1093/sysbio/syy032

23. Bouckaert, Remco R and Heled Joseph (2014) DensiTree: Seeing Trees Through the Forest bioRxiv, 2014, http://dx.doi.org/10.1101/012401

24. Edgar RC (2004) MUSCLE: multiple sequence alignment with high accuracy and high throughput. Nucleic Acids Research 32(5):1792–1797. PMID: 15034147. DOI: 10.1093/nar/gkh340

25. Hedges S, Sagar Baral H, Timmins RJ and Duckworth JW (2008) Bubalus arnee. In: IUCN 2008. 2008 IUCN Red List of Threatened Species. http://www.iucnredlist.org.

26. Hard JJ, Gross M R, Heino M, Hilborn R, Kope R G, Law R and Reynolds J D (2008) SYNTHESIS: Evolutionary consequences of fishing and their implications for salmon. Evolutionary Applications 1:388–408.

27. Devine J A, Wright P J, Pardoe H E, Heino M and Fraser D J (2012) Comparing rates of contemporary evolution in life□history traits for exploited fish stocks. Canadian Journal of Fisheries and Aquatic Sciences 69:1105–1120.

28. Pigeon Gabriel, Marco Festa□Bianchet, David W Coltman, Fanie Pelletier (2016) Intense selective hunting leads to artificial evolution in horn size. Evol Appl. 2016 Apr; 9(4): 521–530.

